# Faster growth with shorter antigens explains a VSG hierarchy during African trypanosome infections: a feint attack by parasites

**DOI:** 10.1101/131029

**Authors:** Dianbo Liu, Luca Albergante, David Horn, Timothy Newman

**Author notes:** These authors contributed equally (David Horn, Timothy Newman).

## Abstract

The parasitic African trypanosome, *Trypanosoma brucei*, evades the adaptive host immune response by a process of antigenic variation that involves the clonal switching of variant surface glycoproteins (VSGs). The VSGs that periodically come to dominate *in vivo* display a hierarchy, but how this hierarchy arises is not well-understood. Combining publicly available genetic data with mathematical modelling, we report a VSG-length-dependent hierarchical timing of clonal VSG dominance in a mouse model, revealing an inverse correlation between VSG length and trypanosome growth-rate. Our analysis indicates that, among parasites switching to new VSGs, those expressing shorter VSGs preferentially accumulate to a detectable level that is sufficient to trigger an effective immune response. Subsequent elimination of faster-growing parasites then allows slower parasites with longer VSGs to accumulate. This interaction between the host and parasite is able by itself to explain the temporal distribution of VSGs observed *in vivo*. Thus, our findings reveal a length-dependent hierarchy that operates during *T. brucei* infection, representing a ‘feint attack’ diversion tactic utilised during infection by these persistent parasites to out-maneuver the host immune system.

**Significance Statement:** The protozoan parasite *Trypanosoma brucei* causes devastating and lethal diseases in humans and livestock. This parasite continuously evades the host adaptive immune response by drawing on a library of variant surface proteins but the mechanisms determining the timing of surface protein – host interactions are not understood. We report a simple mechanism, based on differential growth of parasites with surface proteins of different lengths, which can explain the hierarchy of variants over time. This allows parasites to evade host immune responses for extended timeframes using limited cohorts of surface proteins. We liken this strategy to a military ‘feint attack’, that enhances the parasites ability to evade the host immune response. A similar mechanism may also operate in other important pathogens.

## Introduction

*Trypanosoma brucei*, a species of parasitic protozoan belonging to the Trypanosoma genus, is responsible for African sleeping sickness in humans and nagana in animals. The diseases account for thousands of human deaths per year and pose a serious humanitarian and economic threat to developing countries. The *T. brucei* parasites display variable surface glycoproteins (VSGs) [1-3], which coat the cell surface. VSGs are recognised by the adaptive immune system of the host, which reacts by mounting a VSG-specific immune response. However, each *T. brucei* cell has the ability to dynamically change its specific VSG, therefore forcing the immune system to continuously adapt to the ever-changing ‘antigenic landscape’. This process of immune evasion can continue for years in a single host. Other important parasites that cause malaria and Giardiasis use related mechanisms for immune evasion and persistence.

Given their importance as mediators of a successful infection, the mechanisms that control the switching and expression dynamics of genes encoding VSGs and the resulting host-parasite interactions are active fields of study. Several aspects of the dynamics of VSG expression have been elucidated. For example, it is known that only one of approximately 15 VSGs, found in specific genomic locations close to the telomeres (so-called expression sites), is active in any given cell [4]. However, a significantly larger number of genes (up to two thousand) are present in other genomic locations. The genes found in this archive need to be copied to the active site to be expressed. In this *VSG* gene repertoire of *T. brucei*, less than 15% of the genes are intact. The vast majority are pseudogenes or gene fragments [5, 6] that, nevertheless, can come together to form functional chimeras [7, 8].

Several previous efforts have been made to use mathematical models to understand the complex population dynamics of *T. brucei* and the interaction of the parasites with the host immune system. These models have considered various factors that potentially underlie the unique population behaviours of *T. brucei*, especially the semi-predictable order of appearance of VSGs *in vivo*. The range of mechanisms considered includes different probabilities of activation of VSGs, differential switching rates of variants, density-dependent differentiation from the replicative (slender) form to the non-replicative (stumpy) form of the parasites [2, 9] and clustering of variants [9]. In this article, a fundamental principle underlying antigenic ordering is proposed and validated against *in vivo* experimental data.

Prior observations indicated that parasites expressing VSG-2, which is one of the shortest VSGs of the library (12^th^ percentile) [10], consistently appears in the first-relapse populations *in vivo* [11] and often outcompetes *T. brucei* expressing other VSGs *in vitro* [4, 10, 11]. In addition, VSGs in *T. brucei* have a particularly wide range of lengths. These facts led us to ask whether *T. brucei* clones expressing shorter VSGs could have a survival advantage.

To this end, we constructed a minimal mathematical model to study the potential effect of differing VSG lengths on parasite population dynamics. The model was tested against *in vivo* experimental data obtained from mice [8] and was used to explore how differential VSG-switching, or impact on growth, could contribute to the survival of the parasite in an immuno-competent host. Modelling indicates that faster growth of parasites expressing shorter VSGs extends the duration of infection. This particular model also explains the hierarchy of VSG expression observed *in vivo*.

## Results

### Modelling VSG expression dynamics

One striking aspect of *T. brucei* VSGs is the broad distribution of lengths (Figure 1A) despite those genes apparently having similar functions [4, 10]. This indicates that the length of VSGs may have an intrinsic biological significance. Inspired by previous experimental observations that *T. brucei* variants with shorter VSGs often dominate *in vitro* [4, 10, 11], we used a mathematical model to explore the potential effect of length variation of VSGs on the intra-host parasite population dynamics. In creating a simple mathematical model of *T. brucei* infection focused on the length dependence of VSGs, we introduce three different plausible molecular mechanisms that potentially relate the length of the active VSG and the behaviour of the parasite population as shown in figure 1B: 1) VSG length-dependent switch (DS) in which *T. brucei* with shorter VSGs switch surface antigens less frequently; 2) VSG length-dependent activation (DA) in which shorter VSGs are more likely to be activated; and 3) VSG length-dependent growth (DG) in which parasites with shorter VSGs replicate faster. We also considered a ‘negative control’ null model in which all parasites have the same switching and replication rates, and all VSGs are equally likely to be activated.

Considering a population of *T. brucei* in the bloodstream of the host, we denote the density (number of parasites per ml of blood) of variant *k* by *N*_*k*_ The intrinsic growth rate of each variant depends on its VSG-length. We denote by *r*_*k*_ the length-dependent net growth rate of variant *k* (which accounts for cell replication and cell death due to factors other than adaptive immune killing, and is restricted to the biologically plausible range of 0 to 4 replications per parasite per day). The antigenic switching rate of each variant is also VSG-length-dependent and is donated by α_*k*_ (and is restricted to the biologically plausible range of 0 to 2×10^−4^ switches per parasite per replication). A putative VSG-length-dependent antigenic activation rate of variant *k* is dictated by *Q_k_*, which describes the preferential switching to VSG of variant *k* in the population, and is normalised to unity *I*_*k*_ denotes the rate of adaptive immune killing of variant *k* and is set to the biologically plausible value of 5 per parasite per day [9] and the acquired immune killing threshold is chosen equivalent to the detection of each variant when it reaches ˜ 10^3^ parasites/ml and with a time delay of 4 to 5 days [12]. The specific forms of the length-dependences of *r*_*k*_, α_*k*_, *Q*_*k*_, *I*_*k*_ are described in detail in the Materials and Methods.

Thus, we modelled the dynamics of *T. brucei* parasites in the bloodstream of the host using the following system of equations:

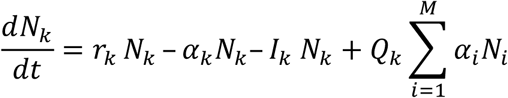

For a given value of *k* (i.e. a given variant) the left-hand-side describes the net change of *N*_*k*_ over time. The right-hand-side comprises the set of biological processes through which *N*_*k*_ can change 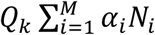 denotes the net switching rate of the whole *T. brucei* population to a specific variant *k*, where *M* is the total number of variants. Note that in our model the immune system adapts to (i.e. recognises) VSG variants over time. In particular, the modelled adaptive immune system will initiate killing of a given VSG-expressing clone after it reaches a set threshold and will continue killing the clones until the infection has been cleared or the host dies (see Materials and Methods for more details). The system of ordinary differential equations was numerically solved to study population dynamics of the parasites in the bloodstream of the host over a 30-day timeframe. All the infections were started with the same initial condition of 1000 parasites (per ml of blood) of a single variant with VSG length = 1500 bp. The distributions of VSG lengths and immune responses to each VSG were tracked over time.

Potential cross-reactivity of the adaptive immune system was not considered in our model. Another feature absent from our model is quorum sensing, balancing proliferative slender-form and arrested stumpy-form parasites. To the best of our knowledge there is no evidence linking quorum sensing to antigen length (Supplementary Figure 1) [13]. In addition, the vast majority of natural infections are characterised by a low level of bloodstream-parasitaemia, in which case quorum-sensing would not be expected to have a major impact.

In the null model, the distribution of VSG lengths and the detected VSG spectrum by the acquired immune system tend to coincide 15 days after infection (Fig. 1C). Similar dynamics are observed in the length-dependent switch (DS) model (Fig. 1D). Interestingly, a different scenario emerges in the length-dependent activation (DA) model: we observe a weak ‘disassociation’ between the expressed VSGs and the VSG spectrum detected by the adaptive immune system (Fig. 1E). A much stronger disassociation is observed with the length-dependent growth (DG) model (Fig. 1F). In this case, we see that the detected VSG spectrum lags significantly behind the expressed VSGs (Fig. 1F). Moreover, the length distribution of the expressed VSG initially moves towards shorter antigens, but by day 15 the trend is reversed and the distribution shifts toward longer VSGs.

This analysis indicates that VSG length-dependent growth rate, and to a lesser extent, VSG length-dependent activation rate, would allow *T. brucei* to establish a more robust infection. This is because the adaptive immune system is unable to generate responses that match the actual distribution of antigens expressed by the parasites in time, due to the inevitable lag in detection by the adaptive immune system. This is reminiscent of a ‘feint attack’ in military tactics. The *T. brucei* population diverts the host-acquired immune system using variants with shorter VSGs, allowing variants with longer VSGs to emerge from below-threshold sub-populations.

### Experimental infection data confirm our theoretical predictions

VSG-seq has been used recently to investigate the dynamics of *T. brucei* infection in an immune-competent mouse model over a period of several weeks [8]. VSG-seq is a variant of RNA-seq to quantitatively track different VSGs and the abundance of cells expressing those VSGs. We used the data from this study to monitor the distribution of VSG lengths over time (see Materials and Methods).

In all four mice studied, the distribution of VSG lengths shifts towards shorter VSGs during the initial phases of infection and then shifts towards longer VSGs in the later phases (Fig. 2 and Supplementary Figures 2-5). This behaviour matches the predictions of our DG model, and to a lesser extent our DA model. Fig. 3 shows both the distribution of VSG length and the VSG length of the clone with the largest number of parasites detected (the ‘dominant clone’) over the period of infection. In all the mice considered we observe a characteristic decrease followed by an increase in both the distribution of VSG length and the VSG length of the dominant clone (Figs. 3*A* and *C*). This behaviour is compatible with both the DA model and the DG model (Figs. 3*B* and *D*). Notably, the DG model allows the parasites to survive longer (white areas) and is therefore the model that provides the greatest advantage to the parasite.

### VSG length-dependent growth enhances *T. brucei* persistence

Our analysis suggests that VSG length-dependent growth would enhance, by itself, persistence. To further explore this aspect, we used our model to further investigate the time needed by the immune system to kill all parasites in an infection (the ‘extinction time’ when all the VSGs available are exhausted) under each of the mechanisms discussed above, excluding biological strategies such as the assembly of VSG mosaics.

A statistically significant difference in the extinction time can be observed across the three models. In particular *T. brucei* survive significantly longer when the DG model is considered (Figs. 4*A* and *B*), while the DA and DS models fail to provide a significant advantage over the null model.

To provide a more precise quantification of this difference, we tracked the time taken by the adaptive immune system to detect all the VSGs under each different mechanism. As illustrated by Figs. 4*C* and 4*D*, the DG mechanism produces a wider distribution of immune detection times, with some VSGs being detected earlier and others being detected later. This wide distribution is not observed in the other models (Figs. 4*C*). This indicates that in the DG model, trypanosomes expressing shorter VSGs are sacrificed at the beginning of the infection, to delay the detection of others, as in a feint attack, thereby increasing persistence of the infection.

## Discussion

*T. brucei* carries a large library of *VSGs* in its genome, which allows a population of parasites to express a broad diversity of antigens, thus limiting the ability of the adaptive immune system to mount a curative immune response. While a diversity of antigens is necessary to evade the immune response, it is equally important that different antigens emerge in a controlled way, in such a way to exploit the limited ability of antigen-presenting cells to identify and expose antigens present at a low concentration. This delays exhaustion of available VSGs and allows the parasite to survive longer. In this article we have explored whether the length distribution of VSGs could be used to provide such a molecular mechanism.

Among the models considered, the hypothesis that the length of the expressed VSG causes differential growth in *T. brucei* was shown to reproduce features of *in vivo* experimental data, and provides significant increase in the persistence of the infection. Moreover, our model supports the idea that molecular stochastic processes can lead to a deterministically structured hierarchy of VSG length emergence times. The findings are consistent with the early appearance *in vivo* [11] and dominance *in vitro* [4, 10, 11] of the short VSG, VSG-2. VSG-length-dependent growth may also explain the 30-year-old observation of coincident multiple activations of VSG-5 (aka. 118) variants, derived through different recombination events [14]. We suggest that switch-frequency, long-debated, may have a relatively minor impact on the appearance of different VSGs.

VSG length-dependent growth is analogous to a ‘feint attack’ tactic of parasites that continually diverts the host immune system. Populations with shorter VSGs grow faster and are detected earlier by the adaptive immune system due to increased exposure. This then allows populations with longer VSGs, which exist at sub-threshold levels, to expand. As shown in Figure 1F, the adaptive immune system lags behind the distribution of VSGs in the population, and is unable to catch up until the library of intact *VSG* genes has been exhausted. At this point, we expect that other mechanisms, such as the assembly of mosaic *VSGs* [7] will be needed to maintain the infection. The suggested mechanism also provides a way for the parasite to produce a hierarchy of VSGs, with shorter antigens emerging early and longer antigens emerging later.

Our observations suggest a multistage, evolutionarily optimized, strategy of *T. brucei* spanning establishment, maintenance and persistence phases of infection. During the first few days of infection, a set of metacyclic *VSGs*, those initially expressed in the fly salivary gland, dominate. This stage is followed by activation of a new set of *VSGs* located in the bloodstream expression sites. Recombination, possibly dependent upon shared homology flanking the active *VSG* [15], then allows the activation of ‘archival’ *VSGs,* which replace ‘old’ *VSGs* in expression sites. Finally, mosaic *VSGs* emerge, often assembled from gene-fragments [7, 8, 15, 16]. We propose that *VSG* length-dependent growth rate plays a key role in extending the timeframe over which each of these groups of *VSGs* are effective.

Given that VSG length has a potentially key role in the interplay between parasites and host, it is reasonable to infer that this particular evolutionary pressure maintains a *T. brucei* VSG library with a wide range of lengths. Notably, it has been suggested that the recombination mechanism that underlies the generation of mosaic *VSGs* may occur within the active expression site [8]. If this is the case, longer VSGs dominating later stages of infections will better facilitate segmental gene conversion by providing potentially longer substrates for homologous recombination, thereby potentially enhancing the generation of mosaics [17].

The length of VSGs may affect the ability of the immune system to recognize invariant antigens on the surface of the parasites. For instance, haptoglobin–hemoglobin receptors mediate heme acquisition in *T. brucei* and are located within the VSG layer on the surface of the parasite [18, 19]. These receptors are recognised by the host’s innate immune system to mediate endocytosis of trypanolytic factor 1 by *T. brucei*. Previous work suggested that haptoglobin–hemoglobin receptors protrude above VSG layers [18]. Therefore, VSG length may also impact access to this and other invariant receptors [20], such as ISG65, currently used as an immunodiagnostic antigen [23].

Elucidation of the molecular mechanisms underpinning the proposed VSG length-dependent growth will require further experimental work. It has been suggested that *T. brucei* has developed cell-cycle check-point mechanisms to monitor VSG protein levels to ensure cell surface protection by VSGs [24]. This feature may contribute to differential growth simply because of the time and energy required to transcribe and translate longer VSGs. Such a hypothesis is attractive because VSG represents the most abundant mRNA and protein in the cell with both present at approximately 10% of total cell-load [3]. Shorter genes have also been reported to produce more abundant mRNAs in *T. brucei* [25, 26], potentially facilitating expression of shorter genes. It is also important to point out that even small differences in the growth rate will cause measurable differences at the population level, due to exponential growth and progression to densities reaching >10^7^/ml of blood in mice [8, 27].

While our analysis suggests that antigen length-dependent growth rate is able, by itself, to explain the observed changes in VSG length over time during an infection of *T. brucei* in mice, we cannot rule out the possibility that the other mechanisms, such as VSG length-dependent antigenic activation, immunosuppression [28] or quorum sensing [2], which may operate in the bloodstream, in adipose [29] or in skin [30], may also play a role.

Our model emphasises the important influence of not only VSG length but also the adaptive immune response on the population dynamics of *T. brucei*. We conclude that VSG length-dependent differential growth is a highly plausible strategy used by *T. brucei* to greatly extend its capacity to evade the host acquired immune response.

## Materials and methods

### Model

The system of ordinary differential equations in our model was numerically solved to study population dynamics of *T. brucei* parasites in the blood stream of a host and the interplay with the host immune system during infection under each of the three hypotheses proposed. For each round of analysis, the initial condition was set as 1000 parasites/ml of a variant with VSG length = 1500bp, because this is the mean length of the VSG library used and a common average starting length reported in experimental datasets [8]. The VSG length distribution in the population was tracked over 30 days post-infection.

A deterministic approach was used to simulate antigenic switching among VSGs and a step function was used to mimic detection via the acquired immune system of the host. In particular, the quantities described by the equation in the main text were modelled as follows:

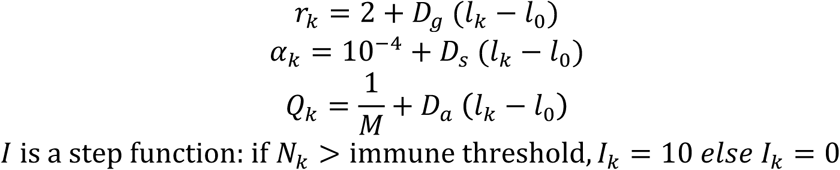

*M* indicates the size of the VSG library and *l*_*0*_ is the median length of all VSGs. The immune threshold used in our study was 10^8^ parasites/ml. The VSG library used for our analysis was kindly provided by the authors of Mugnier *et al.* 2015 [8]. During the initial phases of the infection (1-2 weeks) the direction of the shift of the VSG length distribution depends on the starting VSG clones. Our choice of parameters *D*_*g*_ (4×10^−3^ in DG model)*, D*_*s*_ (2×10^−7^ in DS model), and *D*_*a*_ (proportional to length of each VSG) gives *r*_*k*_, *α*_*k*_ and *Q*_*k*_ ranges of 0˜4 replications/parasite per day, 0-2×10^−4^ switches/parasites per generation and 0-1×10^−2^ respectively for VSGs of different lengths.

*T. brucei* population dynamics data were obtained from published data [8] and additional information was provided by the leading author of the study. In their work, parasites were sampled from the blood stream of four Balb/cByJ mice. The weighted mean of VSG lengths is calculated as the sum of VSG lengths multiplied by their percentages in the population.

## Interplay between parasites and the immune system

The time before extinction of the parasite population in the host and the time of detection of each VSG clone by the immune system were calculated from 500 rounds of numerical integration with random parameters within the ranges given and based on the VSG distribution described above. The Wilcoxon signed-rank test was used to obtain *p*-values of the difference between different models.

## Acknowledgements

We are very grateful to Nina Papavasiliou and Monica Mugnier (Rockefeller University, New York) for sharing their VSG library and expression data. We also thank Md Al Mamun, Sebastian Hutchinson and Sam Palmer for fruitful discussions. D.L. is supported by Wellcome Trust PhD studentship. D.H. is supported by an Investigator Award (100320/Z/12/Z) from the Wellcome Trust. L.A. and T.N. acknowledge support from the Scottish Universities Life Sciences Alliance.

## Author contributions

DL initiated the project, provided original ideas, developed the codes, performed statistical analyses, and wrote the manuscript. LA provided original ideas, supervised the development of the codes, provided guidance in statistical tests, and wrote the manuscript. DH provided original ideas, supervised the project and wrote the manuscript. TN developed the mathematical model, provided original ideas, supervised the project and wrote the manuscript.

---

**Figure 1.**
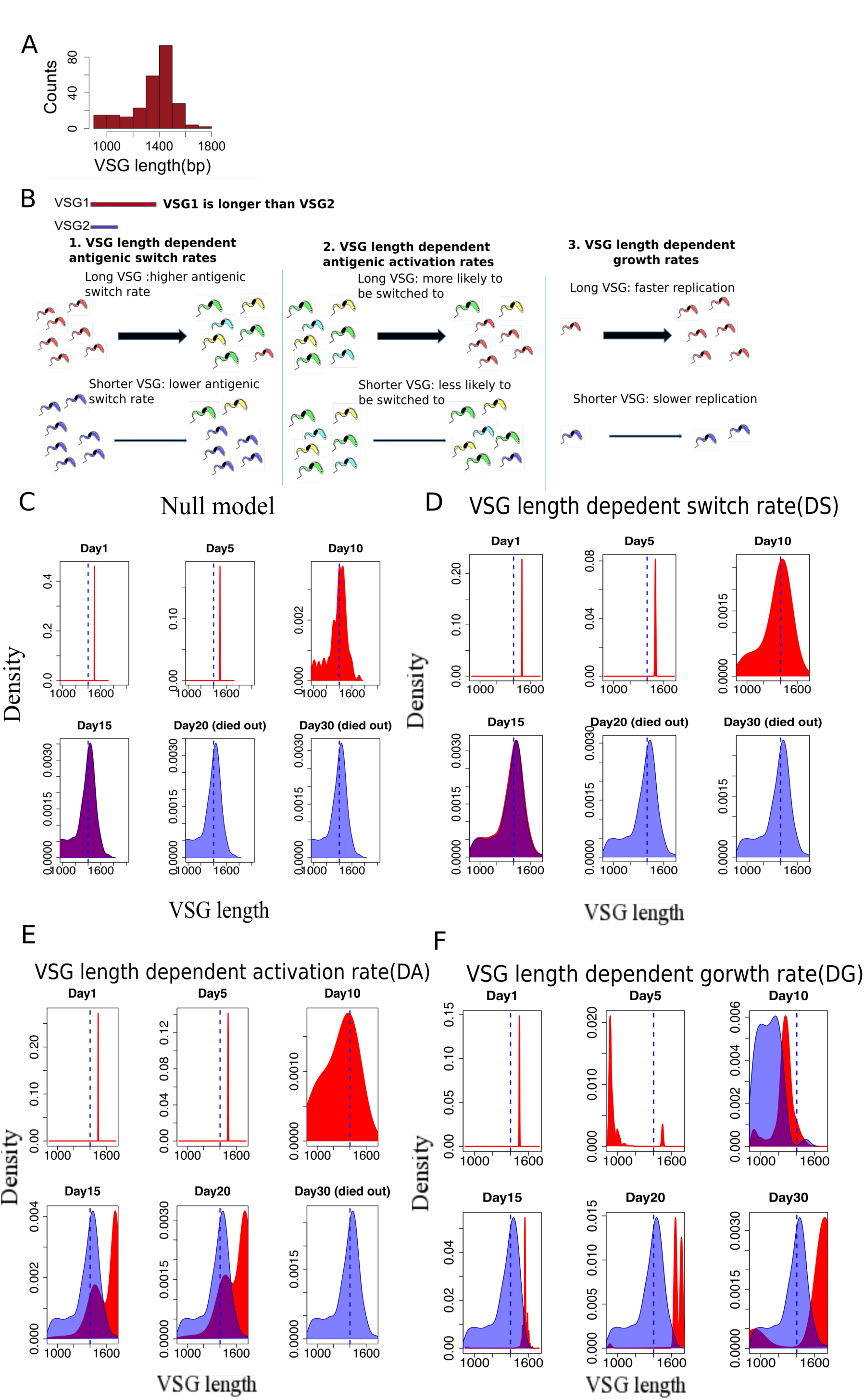
VSG length-dependent population dynamics of *T. brucei*. **. (A)** Distribution of *VSG* lengths [8]. **(B)** Three potential mechanisms are explored in this article: 1) VSG length-dependent switch rate (DS), in which *T. brucei* with short VSGs switch surface antigens less frequently; 2) VSG length-dependent activation rate (DA), in which short VSGs are more likely to be activated; and 3) VSG length-dependent growth rate (DG), in which parasites with shorter VSGs replicate faster. We also considered a ‘negative control’ (null) model in which all the parasites have the same switching and replication rates, and all VSGs are equally likely to be activated. **(C-F)** The red area indicates the expressed *VSG* length distribution in a population of parasites, while the blue area reports VSGs that have been detected by the host adaptive immune system. Blue dashed line indicates the median length of all VSGs in the library. All the densities in figures of this article refer to Kernel density.

**Figure 2.**
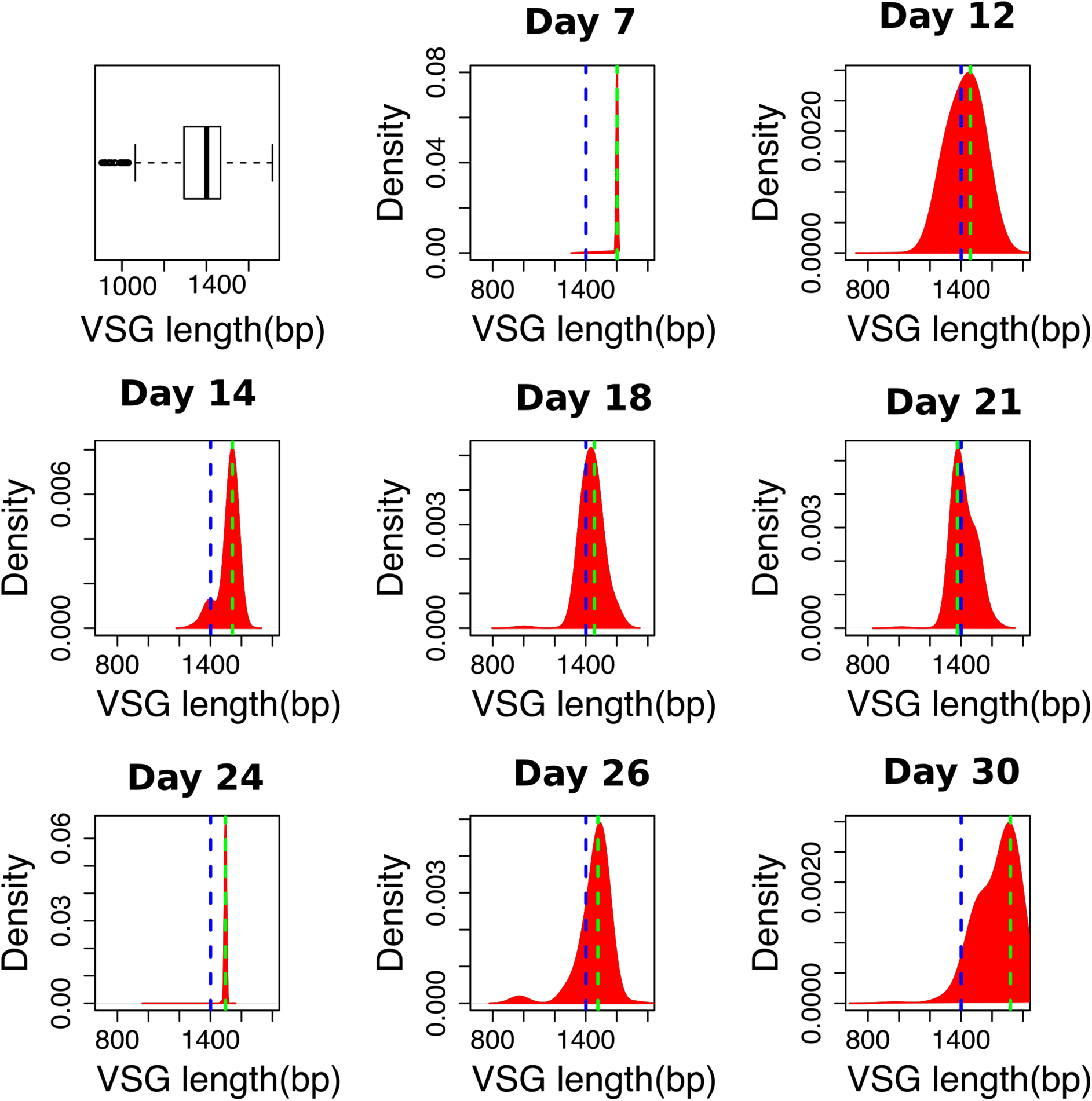
Distribution of expressed VSG length during *T. brucei* infection (Data derived from [8]). The first panel shows the length-distribution of all VSGs in the *T. brucei* genome, while the other panels report the distribution of the VSGs expressed in the bloodstream. Note how the distribution of length shifts towards shorter VSGs during the initial phases of infection and then moves towards longer VSGs in the later phases. The blue dashed line indicates the mean length of VSGs in the genome and the green dashed line is the weighted mean of expressed VSG lengths in the population. This trend was observed in all four mice considered (See Supplementary Figures 2-5 for other mice).

**Figure 3.**
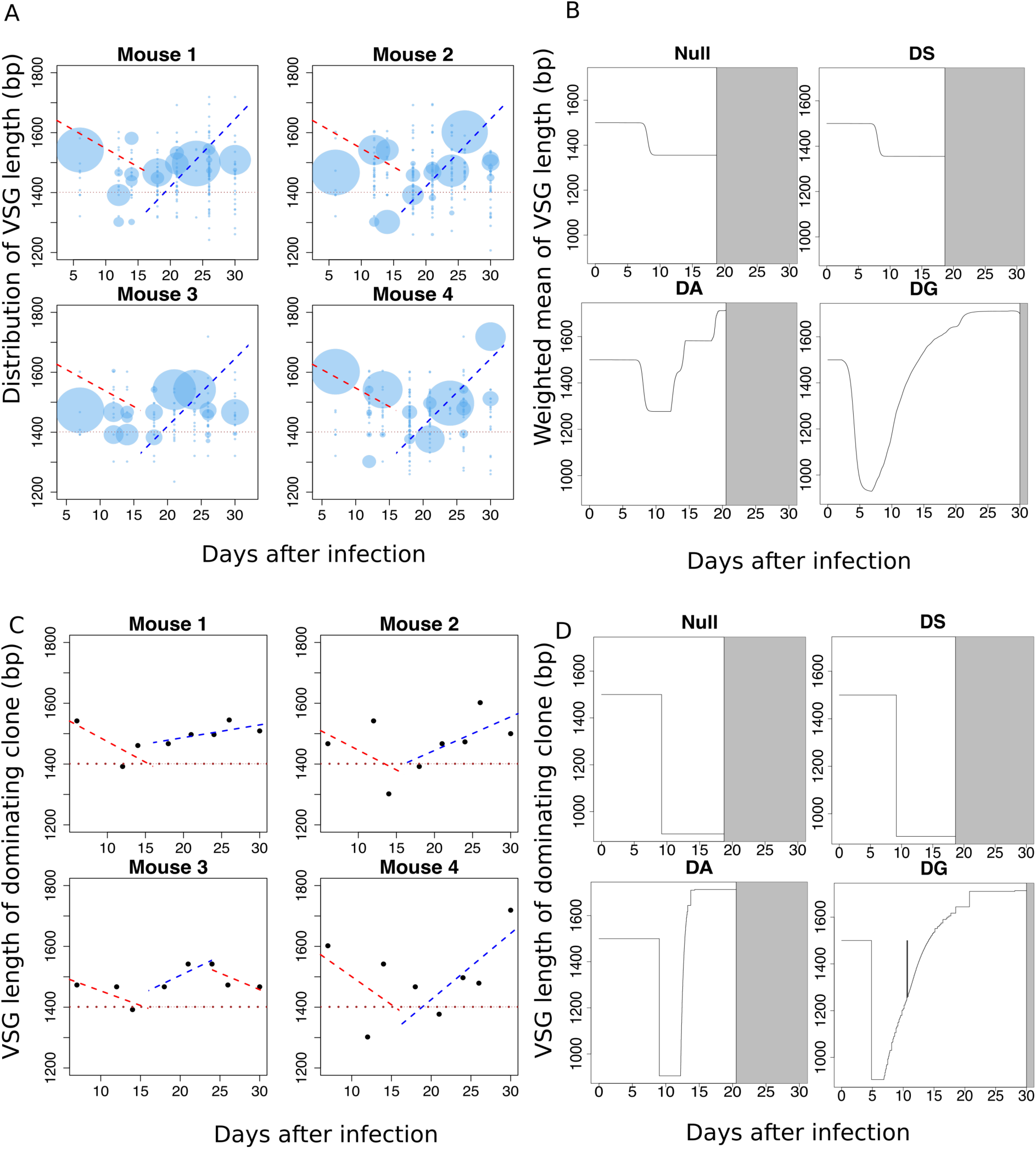
VSG length-dependent growth rate can explain *in vivo* population dynamics. **(A)** The distribution of VSG length in the detectable *T. brucei* population shows a decreasing trend followed by an increasing trend in all 4 mice. The sizes of the circles are proportional to the percentage of VSGs of corresponding length in the population. **(B)** Simulation results show that VSG length-dependent growth rate (DG) is able to reproduce the dynamics of the weighted mean of VSG lengths in experimental data. The grey area indicates the parasite population has died out. The *T. brucei* population survived longest under the VSG length-dependent growth (DG) mechanism. **(C)** The VSG length of the dominating clone shows a decreasing trend followed by an increasing trend in all 4 mice. **(D)** The simulation results for the VSG length of the dominating clone in the population agrees with experimental data and simulation results from panel (B).

**Figure 4.**
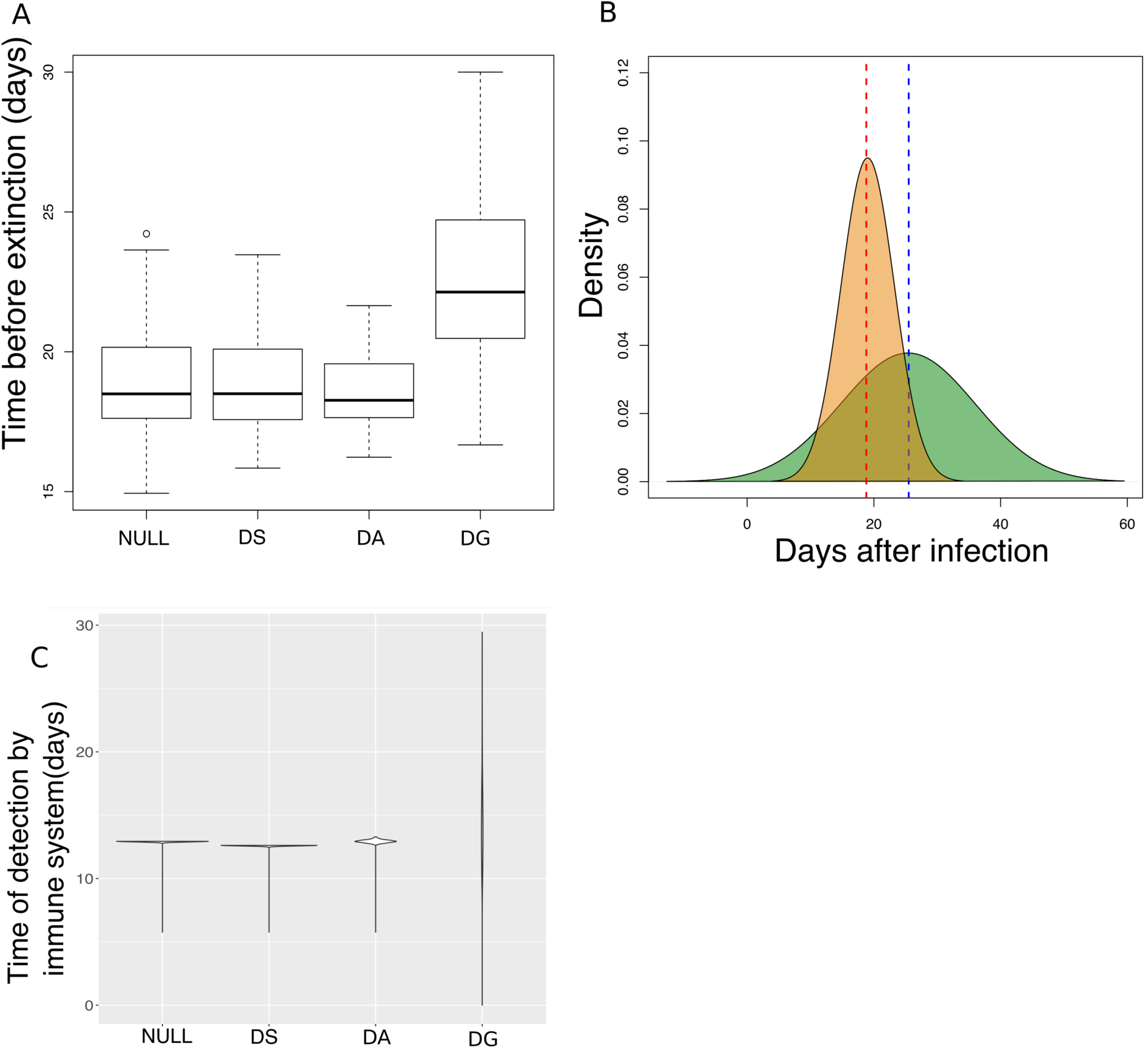
VSG length-dependent growth allows infections to persist for longer (A) The distribution of times before the parasite population died out (extinction) from 500 rounds of simulation with randomly selected biologically plausible parameters. **(B)** The distribution of time before die-out of 500 rounds of simulation of VSG length-dependent growth rate (green) vs. the null model (yellow)**(C)** An example of the time taken by the adaptive immune system to detect each VSG in the population in a simulation. Length-dependent growth gives a wider distribution of immune detection time compared with other models.

